# Measurement of the Topological Dimension of Hippocampal Place Cell Activity

**DOI:** 10.1101/442905

**Authors:** Steven E. Fox, James B. Ranck

## Abstract

The fundamental property of topological dimension of neural activity has never been measured before. We measured the topological dimension of the activity of 89 rat hippocampal place cells recorded during foraging using the neighborhood boundary countdown method. Points were rate vectors forming a manifold in an 89 dimensional rate space Since a boundary has one less dimension than the neighborhood it encloses, reducing the dimension in sequential radial cuts finally arrives at boundary that is two points. The number of cuts required is the topological dimension of the set. Due to sparsity, we used a shell-like boundary with thickness. As expected, we found the *large inductive dimension* of this set of points to be two. To examine the robustness of the method, we used both real and modeled place cells and varied the number of neurons, duration of the time step and smoothing, radius and thickness for the boundary, decrement of the radius with each cut, requirements for the center point, and number of points in the final cluster. With the exception of shell thickness, the result was insensitive to these variations. Knowing the topological dimension allows application of rigorous topological principles to the activity of neuron populations. The method potentially can be applied to other neural classes and other behaviors, to enumerate firing variables when they are unknown, even count the factors influencing neural activity in sleep.

## INTRODUCTION

In mathematics, a *space* is defined by its set of points and its structure. A *topology* is a way in which points in the space can be described collectively. In spaces with the *usual topology* (Euclidean spaces) each member of a set of points in the space has a neighborhood containing a subset of the points. The structure of such a space is in Cartesian coordinates, orthogonal axes of ordered real numbers.

The concept of dimension may seem obvious. It is obvious to all that a line is 1-dimensional; a surface is 2-dimensional; a volume is 3-dimensional; we live in a 3-dimensional world. The concept of topological dimension, which is the subject of this paper, is a generalization of dimension and is not easy or intuitive.

The topological dimension is always an integer; it is *not* fractal. It is the *minimum* number of variables needed to describe the locations of a set of points in a space. Topological dimension is neither a kind of principle component analysis (factor analysis), which approximates the data from variance, nor a kind of multidimensional scaling, which approximates the data from distance between data points. The relationships between data points in a multidimensional space may limit the populated region to a smaller dimensional manifold embedded in that space. Topological dimension describes the number of dimensions of a mathematical space within which the events occur, the embedded manifold. It does not include time and does not describe the path taken between events. The embedded manifold can have many shapes. Topological dimension does not describe the shape. Topological dimension is not an approximation or data reduction. The topological dimension is a characteristic of a topological space and it has the same value in all homeomorphic mappings (1 to 1 and continuous in both directions). Informally, it is the same in all stretching, contracting and twisting of that space. The topological dimension is also the same no matter what metric is used on the space. It is a topological invariant.

Topological dimension of a set of points can be formally defined in three ways (Pears, 1975). It can be defined as the Lebesgue covering dimension, the small inductive dimension or the large inductive dimension. For “well behaved” spaces, (topologically normal) the three definitions are the same. All spaces describing the real world are topologically normal, so for this paper we need not distinguish the different kinds. Here we utilize the large inductive dimension, the only one that can be measured for neural activity data. The other definitions require additional topological knowledge of the space.

The firing of a population of N neurons can be represented in a Euclidean N-space, assigning a rate axis for each neuron. The elements of a vector describing a point within such a neural activity space are the rates of firing of the neurons during a particular time interval, commonly referred to as the “firing rate vector” of the population.

### Why study topological dimension of neural activity space?

To our knowledge no one has ever studied topological dimension of such a neural activity space. The algebraic topological characteristics of persistent homology and cohomology have been used to study data. (e.g.Dabaghian, et al., 2012; Spreemann, et al., 2015; Curto, 2017). None of these studies directly measure topological dimension, although it can be inferred in some cases. Homology and cohomology study geometry or shape, and their uses on data are often focused on the geometric shape of the data. Topological dimension is defined prior to geometry. We do not doubt that neural activity space is to some extent geometric, but there are topological characteristics of neural activity space more fundamental than its geometry, which are defined using topological dimension (e.g. topological embedding and transversality).

While the firing of many neurons is correlated with sensory input or behavior, the firing rates of some neurons have no clear relations with external variables. In cases where there is little overt behavior (e.g. sleep, thinking) we do not even have a first guess of its value, but topological dimension has meaning even when the associated behavior gives no hints. The firing rates of neurons depend not only upon observable parameters, but also upon experimental unobservables that the being may be experiencing. We call the one-dimensional (1D) trajectory of firing rate vectors as a function of time in the rate space the *experiential curve*.

The method we have used here measures the large inductive dimension of a set of points, exploiting the property that a populated subspace with unknown topological dimension n that is contained within an N-dimensional space (n ≤ N) has a *boundary* with nU1 dimensionality. We then take boundaries of boundaries inductively as far as we can. Using this method we measured the topological dimension of a set of points generated by a population of place cells, recorded simultaneously while a rat explored a flat two dimensional surface. Each point is a vector whose components are the firing rates of each of the recorded neurons over a brief time interval. We expected that the topological dimension of the neural activity of many simultaneously recorded place cells of a rat recorded during exploration of a 2D surface would be two.

The large inductive method used here is the same method used by Misner, Thorne and Wheeler (1973)to determine topological dimension in general relativity, (although they do not identity it as the large inductive method). This is an example of the great generality of the concept of topological dimension and of the method for defining and measuring it.

“The *large inductive dimension* of a space X, Ind X, is defined inductively as follows. A space X satisfies Ind X = −1 if and only if X is empty. If n is a non-negative integer, then Ind X ≤n means that for each closed set E and each open set G of X such that E ⊂ G there exists an open set U such that E ⊂ U ⊂ G and Ind bd(U) ≤ n–1 [where bd(U) denotes the boundary of U]. We put Ind X = n if it is true that Ind X ≤ n, but it is not true that Ind X ≤ n - 1. If there exists no integer n for which Ind X ≤ n then we put Ind X = ∞.” (Pears 1975, p. 155)

## MATERIALS AND METHODS

### Theory of the method of large inductive topological dimension

The method we have used to measure the large inductive dimension of the firing of a population of neurons exploits the property that the boundary of a populated subspace with unknown topological dimension (n), embedded in an N-dimensional space, has n-1 dimensionality. A randomly selected point from the set of data points becomes the center of an n-dimensional neighborhood of points with a radius small enough to fit inside the N-space. (The boundary could be any shape; we pick one surrounding a radius for simplicity.) A point on the boundary of that neighborhood is then selected as the center of an (n-1) dimensional neighborhood of points; the boundary of a boundary has dimension n-2. The number of times this process must be applied in order to be left with only the two points at the ends of a curve is an estimate of the large inductive dimension (Ind) of the embedded subspace (manifold) that is populated by the data points. This can be appreciated intuitively in a 3-space that is uniformly and densely populated with points. The 1st radial cut produces a 2D spherical surface, the 2nd, a 1D circle and the 3rd, two points: 3 steps equals topological dimensionality of 3. Optimizing the method to deal with the sparsity of the data points required use of a shell-like boundary with thickness leading to a final pair of clusters of points rather than just two points. Figure 1 illustrates this modified neighborhood boundary countdown method with 2D and 3D sets of points in 3-space.

**Figure 1.**
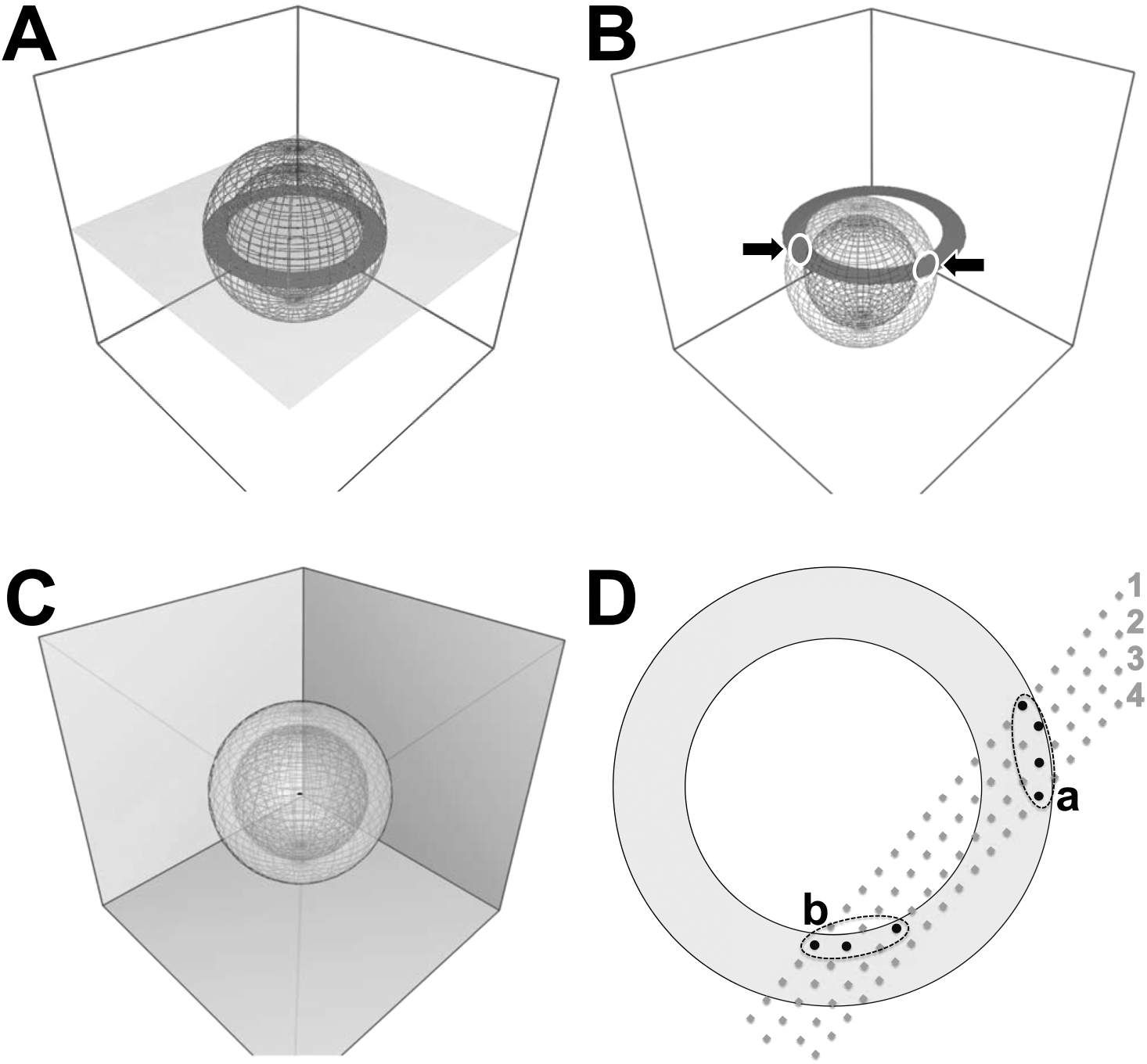
Neighborhood Boundary Countdown. To maximize the number of points included within the boundary of a neighborhood that is an *n-ball*, we give the boundary thickness and call it an (n-1)-*shell,* the contained points describing a rough rather than smooth (n-1)-dimensional surface, so the final result will be two clusters of points instead of two discrete points. **A.** A set of points fills the light grey 2D plane. Cutting with a shell centered at c_1_ leaves the dark grey annulus. **B.** The *second* cut by a shell centered at c_2_ finds two clusters (➔ ➔) along the annulus, a topological dimension of two for the original plane. **C.** The 1^st^ cut of uniformly distributed points in three dimensional space results in a shell that must be cut again to get an annulus and a 3^rd^ time to isolate two clusters, so the topological dimension is three. **D.** Our points are rate vectors in a rate space with dimensionality of the number of simultaneously recorded neurons. Imagine points along adjacent trajectories 1-4 through rate space, cut by the annulus. The initial method included all the points in the grey annulus, so the tangential trajectory, #4, prevented detection of two clusters and required another cut to finish. The new method allows only the first point along a pass through the annulus, so finds only the black points in clusters A and B. For practicality, parts A, B & C are shown in 3-space, but the method generalizes, so the same sets of points (manifolds) embedded in an *n*-space would yield the same values for topological dimension.

In our case the number of neurons recorded is the dimensionality of the embedding space, the rate space. The rate space of N neurons is an N-space having an embedded manifold with a large inductive dimension (n), where n ≤ N, and usually n << N, that represents the number of terms necessary to describe the neuronal activity in the populated subspace.

### How do we initially describe neural activity space to determine dimension?

None of the definitions of topological dimension say anything about time or the path taken between events. Topological dimension describes the mathematical space in which events occur. The events are levels of neural activity measured as the rate of firing of action potentials in a set of neurons. The effect of an action potential outlasts its duration, so we must account for this prolongation in order to count events. We introduce this prolongation with the temporal convolution, using these convolutions as points of the space.

An action potential in a presynaptic neuron may have different consequences in different postsynaptic neurons, and the consequences may be affected by other inputs, so this activity of a neuron may have many terms. However, the form of all these terms is a temporal **<u>convolution</u>**.

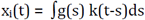
where x_i_(t) is the **<u>activity</u>** as function of time of a single neuron, x_i_, g(s) is usually discontinuous, an action potential at t_0_, i.e. x_i_(t_0_) = 1 and 0 elsewhere, but in neurons without action potentials x_i_(t) is the continuous subthreshold transmembrane potential. k(t-s) is called the **<u>kernel</u>**, is the effect on the postsynaptic neurons at time s after t. The duration and magnitude of this prolonged effect is described by the kernel. A convolution converts a description of a continuous or discontinuous event into a description of continuous activity. The points are the convolutions, not action potentials. We approximate the convolution with a Gaussian smoothing over seconds.

Bittner et al. (2017) report behavioral time scale synaptic plasticity that takes place over several seconds. Schmidt-Hieber and Nolan (2017)reviewed the studies reporting hippocampal pyramidal cell intracellular potential changes lasting several seconds that underlie their place field firing.

We use the average value of this convolution in a time interval as the elements of the points of the rate space. Such a point is commonly referred to as the “firing rate vector” for the population.

This experiential curve plays no role in the topological dimensionality of the space. None of the three definitions of topological dimension say anything about movement along a 1D curve within the space, or a sequence, or include anything about time. For measuring the topological dimension we describe the firing rate space only in terms of the distance between points; time is ignored. Since the firing rates of the population of N neurons define points in a Euclidean space with N orthogonal axes, the distance between two points is given by the generalization of the Pythagorean theorem to N-space. Here, we show the equation for the distance (d_c,b_) in N-space between a randomly chosen center point (c) and some point on the border of its neighborhood (b):

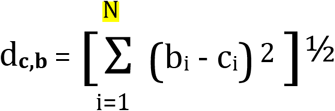

where:

N is the number of neurons, the dimensionality of the rate space

b_i_ is the rate of neuron i at time point **b**, the “border point”

c_i_ is the rate of neuron i at the time point ***c***, the “center point”

So the maximum possible distance between two points in the N-space is the length of an axis (normalized to 100% in our case) times the square root of N (so 943.4% in our case). Rates of activity of 89 different neurons are the elements of the rate vector describing the location of each point in N-space. So the distance between two points c (center) and b (border) in the rate space is the square root of the sum of all element by element differences between their firing rate vectors, where corresponding elements are the rates of the cell in point c and point b. Even if the difference for some cells is zero, the difference for others will contribute to distance between points c and b. For example, in a 4D rate space for cells w, x, y and z, a firing rate vector at a particular time interval (rate of cell w, rate of cell x, rate of cell y, rate of cell z) defines the location of a point in the embedding space. If c = (1,5,3,9) and b = (1,6,3,7), then the distance between c and b is (0^2^+1^2^+0^2^+(U 2)2)^½^ = (5)^½^.

### The neighborhood boundary countdown method for measuring the large inductive dimension

We were provided with recordings of the firing of 89 place cells taken while a rat explored a 1m square open field for one hour, courtesy of Dr. Eva Pastalkova (see crcns.org). Figure 2 illustrates the place field firing rate maps for some of these neurons. These data, only very sparsely populate an 89 dimensional firing rate space. Due to the sparsity of points in the rate space, in order to achieve a correct measure of large inductive dimension it was necessary to use two slightly different radii rather than a single radius to define a spherical *shell* of points lying on a rough, nearly spherical surface, rather than a true, smooth spherical surface, to represent a boundary, and two separated tight clusters of points rather than two points to define a successful estimate of large inductive dimension.

**Figure 2.**
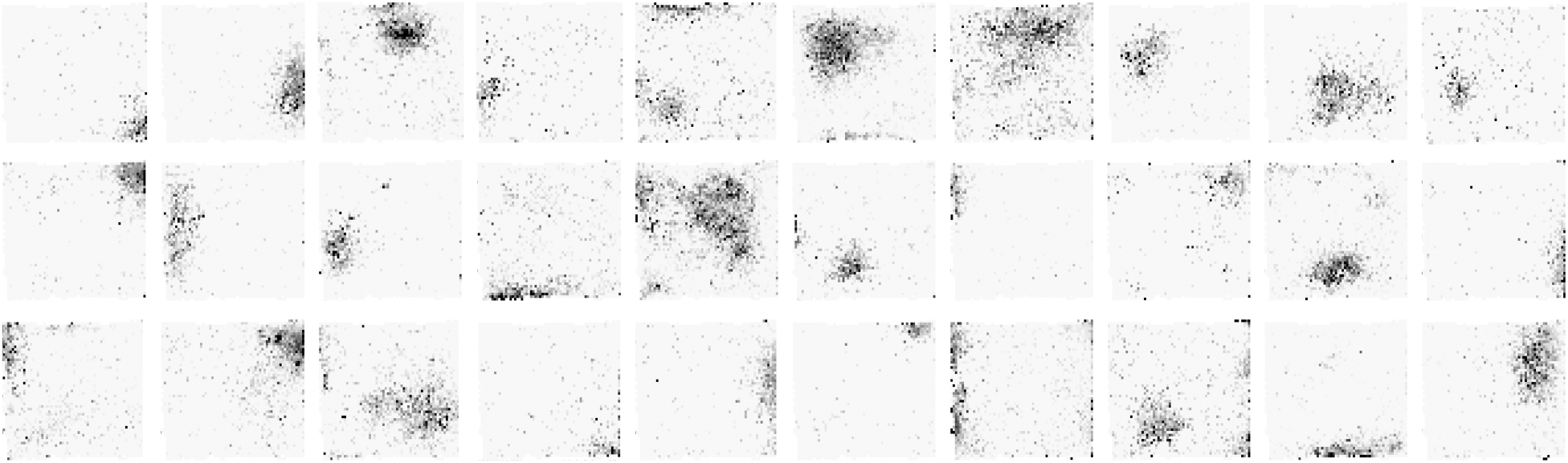
Rate Maps. 30 of the 89 place cells populating the firing rate space. Higher firing rate pixels are darker. The fields vary widely in size and location.

### Controls

Several sets of control data were also used in place of the real place cell data. Artificial place cells were modeled with mean firing rates that were randomly located asymmetric 2D Gaussians on the 1 m square surface either with or without Poisson variation in rate at each time step. We also tested the effects of adding overdispersion-like variation (Fenton & Muller, 1998) in firing rate to these modeled place cells over a ten-second cycle. The analysis was also performed on real and modeled place cell data with time scrambled independently for each cell. Finally, random spike trains were generated such that their distribution of firing rates mimicked the distribution of firing rates in the real place cells. In all cases, the real trajectory of the rat was used for the position data as a function of time.

### Preparation of the raw data

A custom computer program was written to measure the large inductive dimension of both real and modeled place cells and other controls. The algorithm is as follows:

1. For each of the N cells, read the number of spikes per specified time interval into an array.
2. Convolve each cell’s rate array in time with a Gaussian of specified duration, and normalize each cell’s rate to a maximum of 100%, thereby giving the maximum activity of each neuron equal “weight”.
3. Use the rates from step 2 to fill an array representing an N-dimensional space containing a point for each time interval, a “rate vector” whose elements are the temporally smoothed and normalized firing rate of each cell.
4. Set *cut_number* to one.
5. Randomly select an unused candidate “center” point whose elements fit: At least one element greater than a specified low rate and no element greater than a specified high rate. If no such point can be found, go to step 3 (“**No New Point**” failure).
6. Calculate the distances from the selected center point to all axes of the rate space and if the specified radius intersects any axis, go back to step 4 (“**Axis Intersect**” failure, this candidate center point is too close to an axis).
7. Calculate the distances from the selected center point to all other remaining points containing non-zero elements and eliminate from the rate space any points whose distances are not within the specified radius range, thereby forming a spherical shell of points whose (thick) “surface” can be considered to have *cut_number* fewer dimensions than the original populated subspace (neural activity manifold) that was embedded in the N-space.
8. *Optionally* eliminate all but one point along each trajectory (brief time sequence of points) through the radius range, the thickness of the shell.
9. If the number of remaining points is below a selected threshold, go to step 3 (“**Too Few Points**” failure).
10. Calculate the distances between all remaining points
11. **IF:** the remaining points form two tight separated clusters of roughly equal numbers; the topological dimension is equal to *cut_number*; save this measurement of topological dimension (“**Success**”) and continue to step 12. **ELSE:** decrement the radius by the specified amount, increment *cut_number* and go back to step 5 to make a new cut and reduce the dimension of the set of points again.
12. Loop back to step 3 until the specified number of attempts is finished.

To deal with the variance of our successful estimates, we report *measurements* of topological dimension as the modal value of the estimates resulting from thousands of attempts at neighborhood boundary countdown.

## RESULTS

### Successful and unsuccessful trials

The topological dimension is measured as the modal number of countdown steps necessary to arrive at two separate clusters of points. Many different initial radii were tested, usually ranging from 50 to 200% of the axis length in 10% steps. For each of thousands of different randomly chosen initial center points in neural activity space, the algorithm identified all of the points on the boundary of its neighborhood as determined by the test radius. The thickness of the shell defining this neighborhood was usually set to ±5% of the radius. At each step of the countdown process the radius was decreased by some specified percentage, usually 20%, to help reduce the probability of intersecting an axis. For the 89 real place cells and 89 modeled place cells “successful” application of the countdown method generally occurred with radii between 60 and 150% of the axis length. Estimates of topological dimension for random spikes and for real cells with time independently shuffled for each of the cells were strikingly different from those for real cells and modeled cells.

### Real Place Cells

Using this neighborhood boundary countdown method, most attempts to measure topological dimensional of place cell data fail (Figure 3). At small radii most failures result from inadequate numbers of points due to the fact that the rate space is sparsely populated. The occurrence of failures due to insufficient remaining points are called “No New Point” failures and “Too Few Points” failures in the algorithm above. Failure to find a center point that met the required criterion for rates was a “No New Point” failure. When the number of remaining points fell below the limit for the cluster minimum, it generated a “Too Few Points” failure. At large radii most failures are due to intersection of the cut with one of the axes, indicated as an “Axis Intersect” failure in the algorithm. For these place cells, successful estimates of topological dimension were generally less than 10% of attempts. When we allowed only one point per trajectory through the thickness of the shell in order to eliminate failures resulting from our use of a shell as the boundary, successful estimates were generally closer to 1% of attempts.

**Figure 3.**
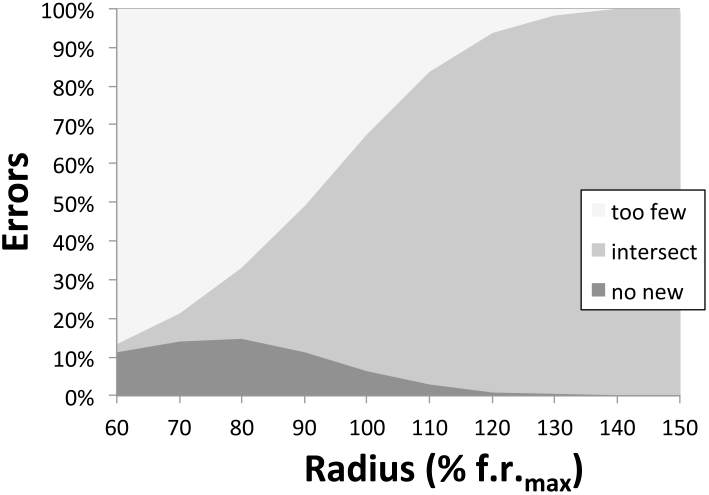
Several Errors Interfere with Dimension Measurements. The rate space is very sparsely populated by the set of points, so most attempts to estimate the dimension of place cell data fail. Small radii find *too few points*. Large radii cut one of the axes (*axis intersect*). Center points had to lie far from the origin, so failures occurred when there was *no new point*. Successful estimates of dimension were usually less than 10% of attempts.

The sparsity of the data in the rate space requires that the “boundary” have thickness (a shell) and a small number of points in the two separate clusters define a success. Because points along a trajectory are generally more closely spaced than points on other parts of the experiential curve, this allows the shell to intersect a segment of the trajectory of the rat that contains multiple points. If a trajectory intersects the shell tangentially, cut twice by the larger, but not cut by the smaller radius, a point in the middle of that segment of the experiential curve may act as the center for the next cut and add an extra dimension to the estimate. Such errors would not occur if it were possible to use a actual spherical surface rather than a shell, and they often led to modal estimate of 3 for the topological dimension of these place cells (Figure 4A). We eliminated such errors by allowing only one point on each trajectory through the thickness of the shell (Figures 1D and 4B). We will refer to the method that allows any point in the shell to be counted as the *All Points* method and the one that eliminates all but one point per trajectory through the thickness of the shell as the *Single Points* method.

**Figure 4.**
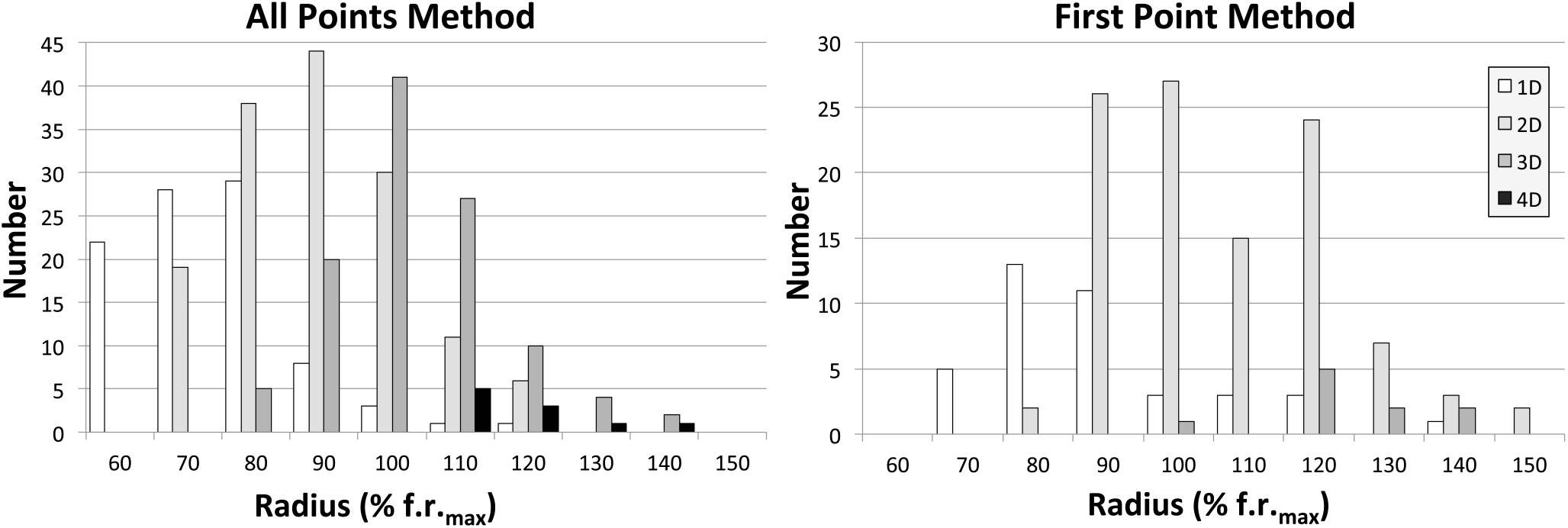
Dimension Estimate Histograms vs. Initial Radius. A. The close points along a trajectory can add an extra dimension (fig 1D) so this graph allowing all of them shows many topological dimension estimates of three. B. Allowing only the first point on a trajectory eliminates most of these 3D estimates. At small cut radii the estimate is often one, likely because successes are limited to cuts through a group of adjacent segments of the experiential curve. As radii increase, the shell can be filled out by points from dissimilar segments. The best estimates occur with the middle range of radii. The largest radii give fewer successes and allow variance to add a dimension.

It is clear in both figure 4A and 4B that only one dimension was found when short radii were used in both the *All Points* and the *Single Points* methods. This too is a result of the sparsity of the data. It is easy to see that for the *All Points* method at short radii a single isolated segment of the experiential curve with closely spaced data points could be cut just once to result in two separate clusters of sequential points after a first cut (c.f. Figure 1D, if trajectories 2 to 4 were absent). For this to occur in the *Single Points* method, requires intersection with several adjacent parallel trajectories through the rate space that are far from other trajectories (c.f. Figure 1D, if trajectories 1 through 4 are all present). As the radius approached the maximum that would fit within the rate space, the shell, defined as a constant percentage of the radius, increased in absolute thickness. This thicker shell allowed the variance in firing rate to populate portions of the shell that would otherwise be empty, so increase by one or more the estimate of topological dimension. Variance could potentially add many dimensions because it can occur in any of the place cells, moving points along any of the 89 axes. But such variance generally has little or no effect on the modal result of our boundary countdown since it increases sparsity thereby increasing the probability of failure. It is clear from figure 4B that, summed across all radii resulting in successful estimates, the modal value for the *Single Points* method is two, representing the two dimensions of the surface the rat explored. But for several radii, the modal value for the *All Points* method is 3 (Figure 4A), representing those two dimensions and another dimension that is actually an error that can be considered to be due to variance and/or an extra cut through the closely spaced points along a tangential experiential curve (Figure 1D).

There are many parameters of the program that are specified before the start of a run. The set that were used most commonly are those of figure 4 (see legend) because they are within a reasonable range considering the velocity of the rat and the distances between points, and they result in a relatively large number of successful estimates for the real place cell data. Some parameters were systematically varied: initial radius, number of cells, minimum points in the clusters, shell thickness, length of time segment, breadth of Gaussian smoothing, decrement in radius per step, and requirements for the center point.

#### Radius

Since the maximum firing rate of each cell was normalized to 100%, the metric of the rate space was percent of maximum rate. The radii that yielded successes ranged from 50% to 200%, most commonly from 60% to 150%. Smaller radii captured too few points and larger ones either intersected an axis or ran out of points, because with the standard smoothing there were no points more than about 250% apart in rate space. As indicated in figure 4 and described above, successful estimates in the smaller range of radii were most likely to give an estimate of 1 for the topological dimension and those in the higher range often added a dimension by inclusion of variance. Short radii are able to intersect the trajectories that passed close to the edge of the floor and some rarely visited regions where the rat might only have moved in one spatial dimension and thereby avoid being filled in by points along the other trajectories that are intersected by larger radii. As you can see in figure 4B and even 4A, the modal estimate of topological dimension was two over a wide range of radii. Collapsing the results for all starting radii, as in figure 5A, makes it clear that the measured topological dimension (the modal value of the estimate) for the *Single Points* method is two, as expected for place cells recorded from a rat exploring a two-dimensional surface.

**Figure 5.**
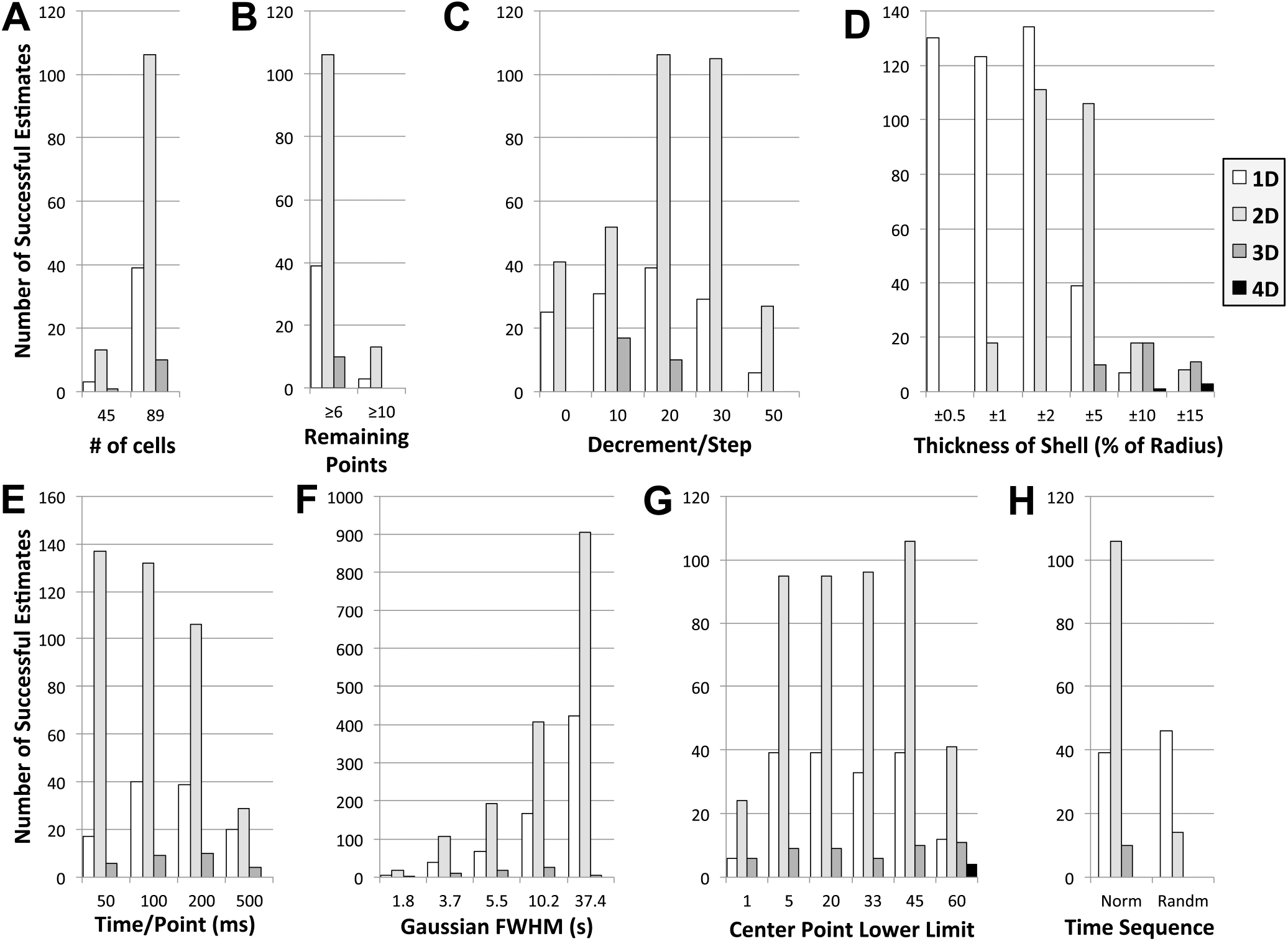
Sum of Successful Estimates Across All Radii. **A**. Using only half of the place cells dramatically reduced the number of successful estimates, but the mode is still two. **B**. Requiring larger numbers of points to be remaining in the final pair of clusters also reduced successes without changing the mode. **C.** When the decrement of the radius after each step was too small there were fewer successful estimates of topological dimension because it increased the likelihood of intersecting an axis. When there was too much decrement too few points were found. **D**. Very thin shells found only 1D, likely due to low probability of including points along multiple trajectories when shell thickness was less than the distance between points along a trajectory. Surprisingly, using thick shells *reduced* the number of successes. **E**. Longer times/point led to fewer successes; shorter ones to more, but computing time was greatly increased. Modal value for topological dimension is still two. **F**. Broader smoothing curves increased success by smearing fields for more overlap; again, computing time was increased. **G.** Requiring at least one element of the center point to have a rate near the middle of its range had little effect on measurement of topological dimension. In fact this lower limit had little effect until below 5% of the maximum rate. **H**. Random spikes yielding a distance distribution similar to real ones produced no successful estimates. We expected that randomizing the time sequence independently for each cell would do the same, but instead it produced dimension estimates of one.

#### Number of cells

In figure 5A, successful estimates of topological dimension at all starting radii are collapsed together for 89 and for 45 real place cells. The number of successful estimates was greatly reduced by reducing the number of cells from 89 to 45, but the measured topological dimension by the *Single Points* method was still two.

#### Points in clusters

It was necessary to allow only 6 remaining points to define two clusters. Those points were required to be distributed nearly equally. If there were only six points they were required to be distributed as either three and three, or two and four in the two clusters. The general rule was that there had to be at least half as many points in the smaller cluster as in the larger one. Figure 5B illustrates two series of runs with thresholds of six and 10 for the minimum number of remaining points. The pattern of results was unchanged, but requiring 10 points resulted in only about one-eighth as many successful estimates of topological dimension.

#### Percent decrement in radius per cut

The length of the radius and absolute thickness for the shell were generally decremented by 20% after each cut to help avoid intersection with the axes. Figure 5C shows that the smaller decrements led to fewer successes, again due to increased intersection with the axes. The 30% decrement produced fewer results of 3, likely by limiting the contributions of variance. At 50% decrement there were many fewer successful estimates due to interception of many fewer points. The modal estimate of topological dimension by the *Single Points* method was still two for all tested decrements of the radius from 0% to 50% per cut.

#### Thickness of the shell

Thickness was specified as a range of plus and minus a percentage of the length of the radius. The most commonly used thickness for the shell was ±5% of the radius, which provided larger numbers of successful estimates of topological dimension than higher percentages without favoring 1D results as occurs with thinner shells. Figure 5D shows that a very thin shell of ±1% or ±2% of the radius resulted in more 1D results, so a smaller fraction of 2D results, likely due to the lower probability of intercepting points along multiple trajectories when the shell thickness was less than the distance between time points along the trajectories. Shells of ±10% and larger resulted in many fewer successful estimates and often included a third dimension, likely due to variance. Large variance may add a dimension by scattering points so they can fill in the gap between what would otherwise be the final two clusters. The reduction in number of successes may have resulted from more axis intersections.

#### Time period per point

For nearly all analyses a 200 ms time step was used to generate points. Time steps from 50 ms to 500 ms were explored and, using the *Single Points* method, resulted in modal topological dimension estimates of two in all cases (Figure 5E). Time steps shorter than 200 ms generated more points, but consumed much more computer time as a result. The 500 ms time step generated fewer points, so resulted in a smaller number of successful estimates than 200 ms.

#### Duration of Gaussian for smoothing

For almost all dimension measurements a Gaussian of width of 3.7 s Full Width at Half Maximum (FWHM) was used. Other widths were explored to show their effect. The longer the duration of the Gaussian used for smoothing the rates of the cells in time (up to 37.4 s FWHM), the larger was the number of successful estimates of topological dimension (Figure 5F). Using a Gaussian with a 3.7 s FWHM yielded many more successes than one of 1.8 s, and seemed to us to be a reasonable duration that wouldn’t smear out place fields too badly. Longer durations were not generally used, but when tested, Gaussian durations of 1.8 s to 37.4 s FWHM gave a similar pattern of results and a modal topological dimension of two.

#### Range of center point

The randomly chosen point that was to become the center point for each cut was generally required to have at least one element with a rate above 45% of maximum. This criterion helped to minimize axis collisions by keeping the selected point away from the origin. Exploration of a wide range of values shows that even 5% was sufficient to produce a very typical pattern of results (Figure 5G). Removing the criterion almost completely by setting the value to 1% substantially reduced the number of successful estimates. The modal estimate of topological dimension by the *Single Points* method was still two for all tested range minima from 1% to 60%.

Randomly scrambling the time sequence for each cell independently did not lead to total failure of the *Single Points* method as was the case for random spikes. Instead, it led to a reduced number of successes and 1D measurements of topological dimension (Figure 5H). This surprising result may be due to less overlap of high firing rates across cells. Since place cells usually fire very slowly and only occasionally fire rapidly, in scrambled data there would be many points composed almost entirely of low firing rates with an occasional single element at high rate. As a result, the smoothing process applied to the randomized sequence of activity would have generated many relatively isolated trajectories in rate space so that two separate clusters of points were found after the first cut.

With the exception of shell thickness, over a wide range of values the above parameters gave a modal estimate of topological dimension of two, as expected. Very thin shells gave estimates of only 1D, likely because intersection with a sufficient number of trajectories to fill the gap between points along a single trajectory never occurred. Very thick shells allowed the variance to contribute at least one additional dimension, as described above. These results for real place cells demonstrate that the method works, is not dependent on a narrow, arbitrarily chosen set of parameters and gives the “correct answer”.

### Controls

Using the *Single Points* method, the modal value of the measurement of topological dimension for the place cell models was also two, with even fewer resulting estimates of one or three than for real cells (Figure 6, note log ordinate). The result was two whether or not Poisson variation in rate at each time step or “overdispersion” (Fenton & Muller, 1998) was included in the model. Using the standard set of parameters, around 20% of the attempts were successful compared to > 2% for real place cells and there was a much smaller fraction of estimates other than two, indicating that the models were much “better” place cells than the real ones. Adding Poisson variance to the modeled cells caused little change in the results. Although not obvious from the log plots, overdispersion increased the fraction of 1D estimates (7.6% for overdispersion vs. 1.5% for no variance), likely by widely scattering the points in rate space. Our implementation of overdispersion did not add another dimension to the data, as might have been expected.

**Figure 6.**
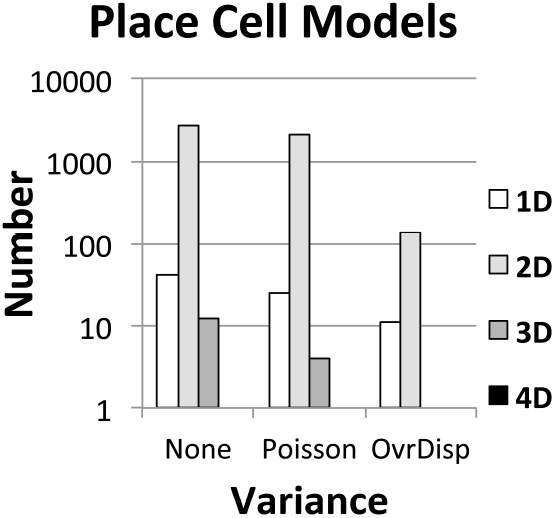
Adding variance to models did not add a dimension. Topological dimension estimates were two for randomly shaped and located 2D Gaussian place cell models both without and with added Poisson or overdispersion-like variance.

The 89 random spike trains that mimicked the distribution of rates in the real place cells did not produce any successful estimates of topological dimension. The expected topological dimension of 89 random spike trains is 89. The sparsity of the data in the rate space made it impossible to reach such a large value due either to interception of small numbers of points, or intersection with an axis with large radii.

## DISCUSSION

Note that the time sequence of points in the rat’s trajectory on the 2D surface is not assigned any special significance; each point is treated as an observation independent of the point that preceded or followed it. However, it seems that in very sparse data sets its relatively high spatial resolution does introduce important consequences.

The activity of these place cells only sparsely populates the rate space. Long-term recording has limited ability to reduce this sparsity, since the data form a manifold of small dimensionality embedded in the many-dimensional rate space. Very small values of the cutting radius find an insufficient number of data points because of this sparseness.

Recording more neurons just increases the number of dimensions without reducing sparsity. Because of the sparsity, as the length of the radius is increased, the successful estimates at the still relatively short radii indicate there is only one topological dimension (see Figure 1). This single dimension represents the “experiential curve”. For perfect place cells without variance or additional cognitive or sensory-behavioral correlates, this curve would correspond to the trajectory of the animal through space-time. Further increases in radius and sigma identify more topological dimensions as they intersect broader ranges of data points from multiple segments of the experiential curve. Large variance can scatter data points along trajectories and across repetitions to further increase the measured number of topological dimensions.

So there are two practical limitations on the topological dimensionality measurements for neural activity: sparsity and variance. Because of sparseness, the method sometimes (especially at short radii) finds the final two separate clusters of points, the ends of the one dimensional line segment that terminates the countdown, by cutting the one-dimensional experiential curve. **This occurs because that sequential points along a short segment of the experiential curve tend to be much closer to each other than they are to the points along a different segment.** Highly variable rates allow erroneous filling in of the rate space with points scattered by the large variance, in what would otherwise be empty space between a terminal pair of clusters. This error occurs more commonly as the thickness of the shell increases. With large values of variance and sigma it can even add more than one dimension to the estimate (see Figure 5D).

To deal with sparsity, the inductive method was modified to use a shell for the boundary, rather than a surface. Our first analyses, accepting all points contained in the shell, seemed to indicate that the topological dimension was three, not the two that was expected for the 2D floor. Because the shell has thickness, the extra dimension is the “experiential curve” itself, the activity along the rat’s path. In the same way that the closely spaced points along the curve indicate one dimension with very short radii, they can add a dimension with larger radii. We have now eliminated the extra dimension by allowing inclusion of only one intercepted point in each pass through the thickness of the shell (see Figures 1D and 3). The modal estimate was two for both the real place cells and models of place cells both with and without variance, and it was not sensitive to exact radius or thickness of the shell. The modal estimate was unchanged by reducing N to 45, but N=30 was insufficient to ever succeed. Applying the method to the same number of pseudo-randomly distributed points in a rate space of the same dimension achieved no successful estimates due to the extreme sparsity of the data.

A topological dimension is always an integer. Informally and intuitively we can say that a topological dimension is the number of “ways” that a variable at a point can change continuously. A topological dimension is the minimum number of terms necessary to describe a set of points in a space. The number of ways a curve can change can be described not just in numbers, but also in words or figures. Perpetual systems are continuously active systems such as those controlling blood pressure, body temperature, osmolarity, blood concentrations of some hormones, length of a skeletal muscle, tension of a skeletal muscle, etc. The activity of each these systems is one dimensional because the activity of **<u>all</u>** of the neurons change in only one way. Although they may have different thresholds and gains, when they change, all of the neurons change together. A possible confusion: informally, a topological dimension describes the change in neural activity space, not whether it is increasing or decreasing. The same subset of neurons increasing their rate by any amount or decreasing their rate by any amount has the same topological dimension.

The topological dimension does not care about excitation or inhibition.

Whereas the conclusion was predictable for place cells because the firing correlates are well known, there are many neuronal classes and conditions for which there are an unknown number of terms necessary to account for the firing, e.g. firing with no change in behavior, as in sleep. The topological dimension of these firing rate manifolds is useful because: 1) it allows more extensive mathematical analyses to be used, e.g. maps between manifolds, combining manifolds, and 2) it allows us to define circumstances in which the experiential curve intersects itself, i.e. in which something is the “same”, as is required in learning and recognition.

## ACKNOWLEDGEMENTS

We are greatly indebted to Dr. Eva Pastalkova who provided us with files containing her recordings from 89 simultaneously recorded place cells. Dr. John Kubie provided rate map visualization software, as well as useful discussions during the development of this method.

